# Predictability failure in glucose-insulin system for ICU patients

**DOI:** 10.64898/2026.08.26.747449

**Authors:** Deepjyoti Ghosh

## Abstract

Modern medicine implicitly assumes that physiological responses to intervention are predictably determined by administered treatments. However, physiological systems containing intrinsic delays between the detection of a stimulus and the biological response may violate this assumption. We investigate the human glucose–insulin system as described by the Ultradian model and mathematically demonstrate that clinically relevant forcing protocols—such as pulsatile insulin delivery and step-wise glucose infusion, both commonly used in intensive care units (ICUs)—can induce sustained temporal chaos that may hamper accurate prediction of the physiological response. If not accounted for, these chaotic dynamics could create difficulties in achieving optimal dosing and timing when administering glucose and insulin in clinical or home care settings. This phenomenon, termed delay-induced uncertainty (DIU), arises from the interaction between physiological delay, intrinsic shear near a limit cycle, and external forcing. Using the Ultradian glucose–insulin model, we compute top Lyapunov exponents to quantify predictability. Across a range of pulsatile and step-wise forcing regimes, including stochastic amplitudes drawn from Markov processes, we observe positive Lyapunov exponents, indicating sustained chaos. Our results suggest that delayed endocrine regulation may fundamentally limit the predictive value of the models used to develop glycemic management strategies, with implications for clinical protocols in the ICU.

## 1 Introduction

Accurate prediction of various natural systems is a key motivator of scientific and mathematical investigation, often for pragmatic reasons: effectively modeling the behavior of dynamic systems such as weather patterns, volcanic activity, and epidemic diseases has the potential to mitigate risk and save lives. However, mathematical research has shown that such systems often exhibit chaotic behaviour, making them difficult to interpret and thus posing challenges for accurate prediction of outcomes. Substantial theory has been developed to describe how this unpredictability can affect human management of large-scale natural phenomena such as climatic systems or wildlife populations. In contrast, few researchers have explored how the chaotic behavior seen in many models of human physiological systems^1^ could interact with medical treatment. This question may have substantial clinical relevance. The fundamental assumption underlying medical intervention is that it will produce predictable and desirable outcomes, but this predictability may be undermined by the presence of chaotic behavior in the homeostatic systems that regulate human health.

One important feature of biological systems that may lead to chaotic effects is physiological delay. “Delay” in this context does not imply pathology or malfunction, but refers to the inevitable interval between the detection of an input (such as light, physical perturbation, or chemical cues) and the generation of a response (such as physical movement, hormone secretion, or altered gene expression). Building on rigorous theoretical studies of non-uniform hyperbolicity in mathematical dynamical systems, Karamched and collaborators^2^ have identified a novel route through which sustained temporal chaos can emerge in a homeostatic physiological system—specifically, the human glucose-insulin system. This crucial metabolic regulatory system functions as a homeostatic oscillator: insulin secretion by the pancreas increases in response to high blood glucose levels, triggering glucose uptake by cells, and decreases when blood glucose drops. Both excesses (hyperglycemia) and deficits (hypoglycemia) of glucose can have serious health repercussions. Karamched et al. show that the delay between external forcing of glucose (i.e., the addition of glucose from outside the system) and the resulting shift in insulin secretion is sufficient to generate chaotic behaviour, which they term *delay-induced uncertainty* (DIU).

The presence of DIU suggests that the dynamics of the glucose-insulin system may not be fully predictable even in principle. This finding could have important implications for patient care. Glycemic management—the maintenance of a patient’s blood glucose levels within safe boundaries—is a critical component of treatment in the intensive care unit (ICU)^3,4^, and chaotic behaviour could interfere with accurate prediction of how patients will respond to glycemic interventions. Accurate modeling of DIU may therefore shed light on known challenges in clinical management of blood sugar.

Karamched et al model DIU within an idealized finite-dimensional stationary model of the glucose-insulin system known as the Ultradian model^2,5–8^, which they perturb with regular glucose pulses of constant amplitude. This approach provides a compelling demonstration of how chaotic behavior can in principle emerge within this system. However, it does not account for several important features of real-world glycemic management, potentially limiting its clinical applicability. Here, we extend the original DIU framework to incorporate those features:

1. External administration of insulin
2. Non-uniform glucose intake
3. A continuous stepwise flow of glucose mimicking intravenous administration in the intensive care unit (ICU).

Using the *Lyapunov exponent* (see section 4) as our central metric, we examine the presence of chaos in the Ultradian system when subjected to our more realistic model of external insulin and glucose forcing. A positive magnitude for the top Lyapunov exponent signifies the presence of chaos in the system; a negative top Lyapunov exponent indicates the absence of chaos. Under this framework, we show that the dynamics of insulin and glucose administration in the ICU likely amplify the tendency toward chaotic behavior described by Karamched et al.

We discuss relevant research and related theoretical work in Section 2. The Ultradian model used for computation is discussed in Section 3. In Section 4 we explain and define the Lyapunov exponent. To investigate the effect of pulsatile kicks (discrete, instantaneous external forcing events) for glucose and insulin, we first conduct simulations in idealized settings in Section 5 . We kick plasma insulin under an external forcing function that models delta kicks with either constant or randomly drawn amplitudes. For kicking glucose with pulsatile kicks, we choose the kick amplitudes from a Markov process. Further, we develop a step-wise external forcing function in Section 6 to mimic protocols followed in intensive care units (ICU). Finally, we present our conclusions and discuss future directions in Section 7.

## 2 Related work

The direct precursor of our present work is the 2021 paper^2^ from Karamched et al. describing how DIU emerges in the oscillatory behavior of the human glucose-insulin system. These findings emphasize that the presence of delay in physiological systems can produce unpredictable outcomes. The authors combine the Ultradian model of glucose homeostasis with idealized external pulsatile forcing functions that simulate periodic glucose intake from food. They develop simulations to model DIU and explain its impacts using the theory of rank one maps from smooth dynamical systems. The core Karamched framework describes three intrinsic characteristics of the glucose-insulin system that combine to produce chaotic behaviour:

1. **Delay-induced excitability**. The physiological delay between changes in glucose levels and corresponding changes in insulin production produces a weakly stable limit cycle via a supercritical Hopf bifurcation.
2. **Intrinsic shear**. In the context of an oscillatory dynamical system, shear is a measure of the angular velocity gradients near the limit cycle. Karamched’s model indicates that the glucose-insulin system exhibits shear near the limit cycle even when the system is unperturbed.
3. **External forcing**. When a dynamical system receives additional force from outside (in this context, glucose or insulin from an external source), this force interacts with shear to stretch and fold the phase space. This stretching and folding produces sustained temporal chaos.

Karamched et al. subsequently applied the DIU framework to the context of meal ingestion, examining whether delay-induced chaos degrades the capacity of the glucose-insulin system to maintain homeostasis, and explored the implications for obesity and type-2 diabetes mellitus^9^. That study operates in a different parameter regime from the intensive-care setting treated in their original work, whereas the forcing protocols we develop here return to the clinical regime and extend it to administration of insulin and to continuous rather than instantaneous glucose delivery. In this framework, DIU emerges in the glucose-insulin system through a process similar to the linear shear flow model explored by Zaslavsky^10^ and Lin and Young^11^. Several publications by Wang^12–14^ offer helpful context for understanding the dynamic profile of DIU and the development of strange attractors from the theory of rank one maps. Ott^15^ and Wang^16^ additionally provide detailed analyses of perturbing limit cycles for nonlinear systems.

Researchers have developed a wide variety of models to describe the behavior of the glucose-insulin system^17–19^and predict the effects of insulin treatment^8,20,21^. We follow Karamched in basing our experiments on the Ultradian model^5–8,22^(see section 3), which is simple to interpret and has been shown to accurately predict glucose-insulin dynamics in humans.

## 3 Ultradian model

The Ultradian model^2,5–8^is widely used and generally regarded as the simplest physiological model of the oscillatory behavior of the glucose-insulin system. This compartment model has three state variables, viz. plasma glucose *G*, plasma insulin *I*_*p*_ and remote or interstitial insulin *I*_*i*_. The natural physiological delay in the glucose-insulin system is expressed through a three stage linear delay filter modulating the interactions between these state variables. Figure 1 shows the schematic of the Ultradian model, which is described by the following six nonlinear ordinary differential equations:

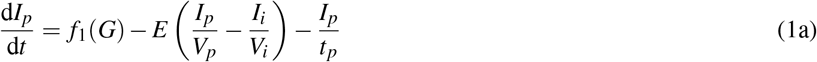

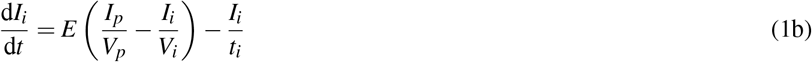

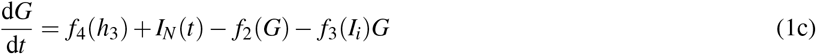

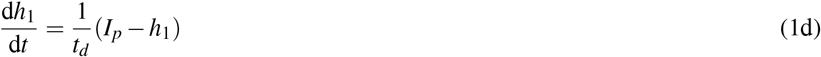

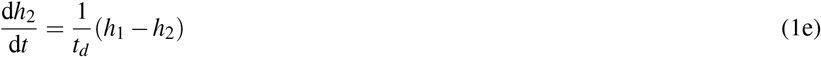

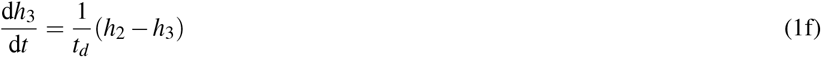

**Figure 1.**
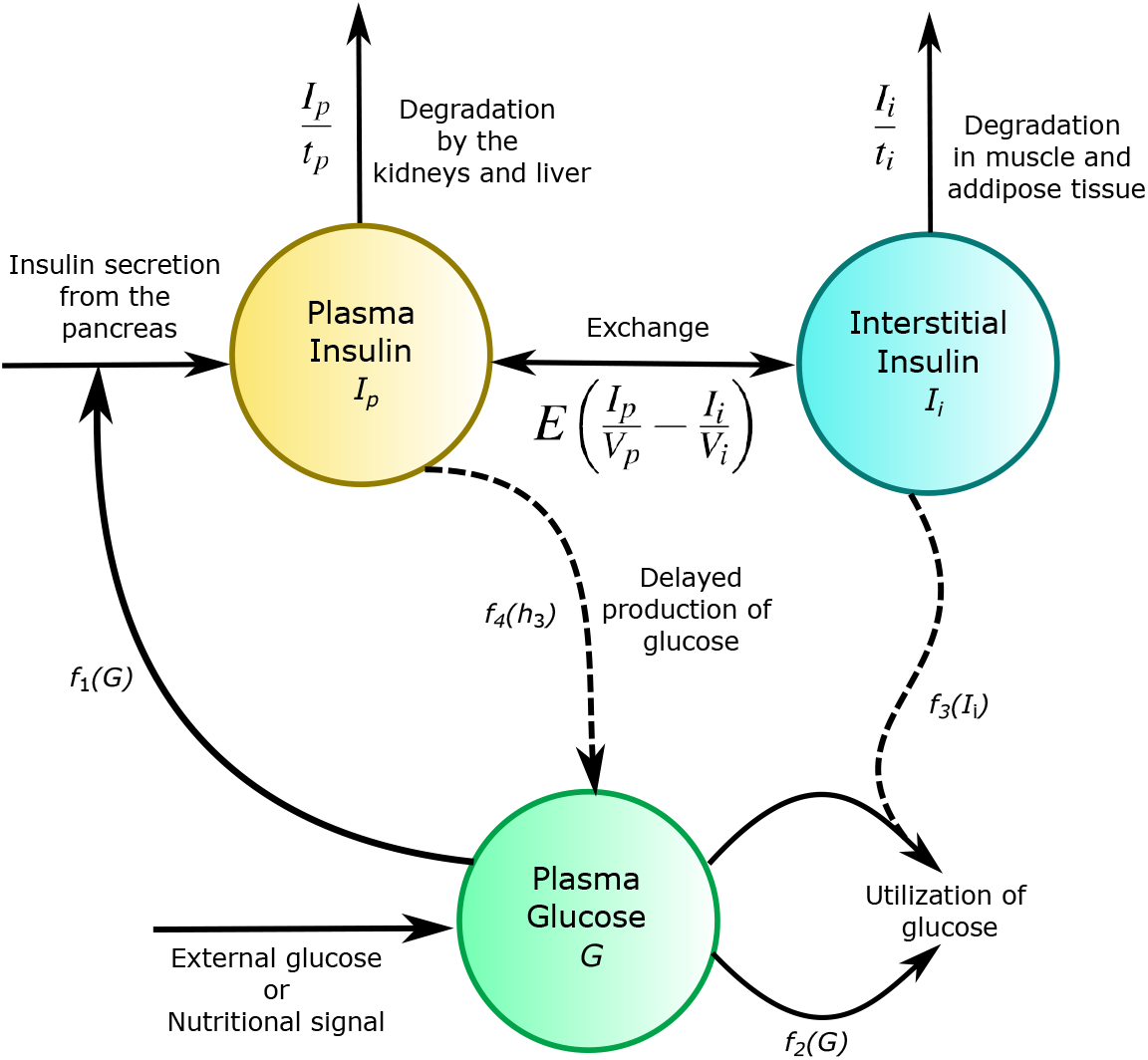
Schematic of the Ultradian model. Diagram illustrating the Ultradian model, including the three state variables, their inputs and outputs, and their interactions.

This model, which we will refer to below as System 1, assumes constant blood and interstitial volumes. The six equations above are derived as follows:

- *f*_1_(*G*) represents the plasma insulin production rate dependent on plasma glucose, where *G* is the total amount of glucose. *V*_*p*_ is the plasma volume and *V*_*i*_ is the interstitial insulin volume. The amount of plasma insulin is represented by *I*_*p*_ and *I*_*i*_ represents the interstitial insulin amount. The exchange between the plasma and interstitial insulin is expressed as their concentration difference along with the rate constant *E* as: 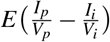 Thus, we arrive at equation 1a.
- In addition to the exchange between plasma and interstitial insulin described in the previous point, interstitial insulin also undergoes degradation in the body through muscles and tissues at a rate 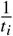, which gives us the equation 1b.
- The plasma glucose level is indirectly determined by the plasma insulin, which is given by *f*_4_(*h*_3_), where *f*_4_ is the delayed production rate from plasma insulin and *h*_3_ is the output of the delay filter. The external forcing drive is represented by *I*_*N*_(*t*). Plasma glucose can be utilized independently of insulin or with interstitial insulin dependence. In equation 1c, *f*_2_(*G*) represents insulin-independent degradation of glucose and *f*_3_(*I*_*i*_)*G* represents insulin-dependent glucose degradation.
- Equations 1d-1f are the three-stage delay linear filters with *t*_*d*_ as the delay parameter. The filter takes *I*_*p*_ as the input and produces *h*_3_ as the output.

The functional forms of *f*_1_, *f*_2_, *f*_3_ and *f*_4_ are represented as follows:

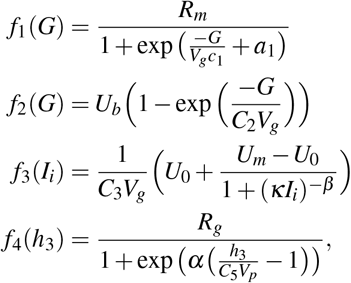

With

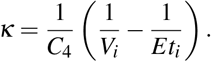

Note that both plasma and interstitial insulin influence plasma glucose levels, though in different ways. The time delay between the production of glucose and its utilization gives rise to glucose-insulin oscillations; here, we are interested in exploring clinical settings in which these delay-driven effects can produce temporal chaos (DIU).

## 4 Lyapunov exponent

Our hypothesis is that DIU is generated in the glucose-insulin system when there is shear present and glucose or insulin are introduced through repeated external forcing. We cannot prove this rigorously, so we use the *Lyapunov exponent* as a mathematical proxy to show the presence of sustained temporal chaos.

The Lyapunov exponent quantifies the rate of separation of two trajectories that were initially very close to one another. Wilkinson^23^ defined the Lyapunov exponent precisely in 2015: Let *f* : *M → M* be a *C*^1^ map on a compact *d*-dimensional manifold *M*. For each *x ∈ M* we denote the derivative of *f* at *x* by *D*_*x*_ *f* : *T*_*x*_*M → T*_*f* (*x*)_*M*, where *T*_*x*_*M* is the set of tangents at the point *x*. Let *D f* denote the derivative cocycle. Then Λ is a Lyapunov exponent for *D f* at *x ∈ M* if there exists *v ∈ T*_*x*_*M* such that:

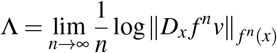

For various forcing signals *I*_*N*_(*t*), we compute the top Lyapunov exponent Λ_max_ for the time-*T* maps induced by the Ultradian model. A positive top Lyapunov exponent, where Λ_max_ *>* 0, indicates the presence of sustained temporal chaos, while Λ_max_ *<* 0 indicates the absence of chaos.

## 5 Modeling pulsatile kicks for insulin and glucose

Expanding on the demonstration by Karamched et al.^2^ that DIU emerges in the glucose-insulin system when it is subjected to idealized pulsatile glucose forcing functions, we provide computational evidence of similar DIU emergence under a more realistic regime of external forcing for both glucose and insulin. For the forcing functions in our experiments we will be using both pulsatile forcing kicks and piece-wise continuous time forcing. These regimes simulate two distinct modes of glucose or insulin administration: discrete consumption/injection and continuous administration via intravenous drip, respectively. For code availability please refer to our Github repository.

We will first consider the Ultradian model using pulsatile forcing functions (Figure 2a) as the external forcing signals for both plasma insulin and glucose. The following equation describes pulsatile kick behaviors mathematically:

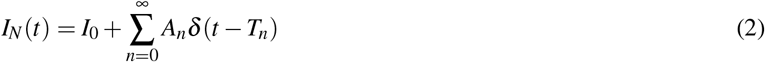

**Figure 2.**
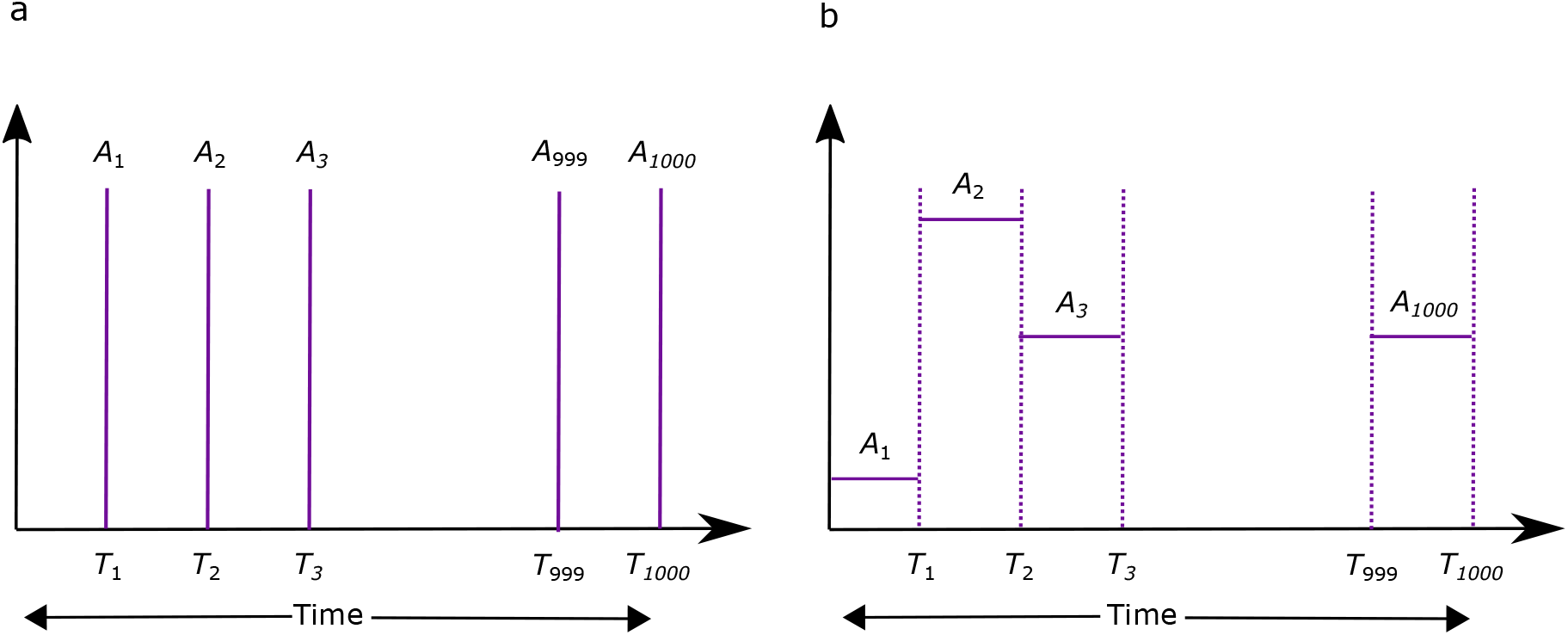
Pulsatile kicks and step-wise continuous forcing. **(a)** Pulsatile kicks, with forcing amplitude *A*_*i*_, applied at each time point *T*_*n*_ with a relaxation period between kicks. For our experiments with pulsatile kicks, *A*_*i*_’s are either constant or randomly drawn from a distribution **(b)** Step-wise continuous forcing, with forcing amplitude held constant between glucose time change points. For our experiments with step-wise forcing, *A*_*i*_’s are randomly drawn from a Markov process.

Computationally, we solve the system 1 using ode23s in MATLAB. We want to examine the presence of DIU when a state variable (plasma insulin or plasma glucose) in system 1 is given an external kick of the pulsatile form described above (Equation 2). For instance, in System 1, *I*_*N*_ represents an external forcing function for the glucose state variable. Between any two kick times *T*_*n*_ *< t < T*_*n*+1_, the ode solver solves the Ultradian model with *I*_*N*_(*t*) = *I*_0_. At the kick time *T*_*n*_, the differential equation solver stops and the glucose state variable receives an instantaneous amplitude change by *A*_*n*_, viz. *G ↦G* + *A*_*n*_. In general, *A*_*n*_ can denote the forcing amplitudes for either plasma insulin or glucose, depending on which state variable is being affected by the external force. For all of our simulations, we take *I*_0_ = 0.

Here we describe how we compute the top Lyapunov exponent Λ_max_ for different lengths of the interval between kicks(we refer to this value as the “inter-kick time”). We vary the inter-kick time from 1 to 100 minutes in increments of 1 minute. Beginning with two solutions of System 1 separated initially by a distance of *d*_0_ = 10^−8^, we apply an external kick at *T*_*n*_ and then compute the separation distance between the two solution trajectories for the state variable. Let this distance of separation be *d*_1_. Then we compute the quantity 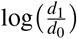 and save it as a vector element. We now renormalize the positions of the two particles so that they are *d*_0_ distance apart and allow a relaxation period equal to the inter-kick time before applying the next kick. This process is iterated 1000 times—giving us a vector of 1000 elements of the quantity 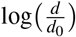, where *d* = *d*_1_, *d*_2_, .., *d*_1000_—for each value of the inter-kick time. For a given inter-kick time, the numerical value obtained by averaging over the 1000 values gives us the top Lyapunov exponent Λ_max_. If the top Lyapunov exponent Λ_max_ is positive it will signify the presence of DIU in the system or vice-versa.

We consider periodic and Poissonian kick profiles. In the periodic case, we have *T*_*n*_ = *nT* for all *n*, where *T* is the inter-kick time. In the Poissonian case, we draw the inter-kick times independently from an exponential distribution with mean *T* . In real life, meal consumption times are not strictly periodic; using Poissonian kick profiles for glucose forcing accounts for this time variability.

The numerical values we used in our experiments for the Ultradian model described in system 1 are in supplement. For more information about this parameter set, please refer to Albers (2017)^22^.

### 5.1 Pulsatile kicks for insulin

As described above, we first examined the case of a uniform insulin kick regime, similar to the simplified glucose kicks in the Karamched model. This analysis will simulate the infusion of insulin for therapeutic purposes and serve as a test of the robustness of Karamched’s predictions of DIU in the Ultradian system. We can computationally verify how the system responds to external forcing for different state variables.

Applying external pulsatile forcing functions to plasma insulin, we compute the top Lyapunov exponent Λ_max_ for inter-kick time values ranging from 1 to 100 minutes. For each inter-kick time, we iterate over 1000 kick relaxation periods, with each relaxation period followed by a kick (given by *I*_*p*_ *↦I*_*p*_ + *A*_*n*_) that instantaneously increases the plasma insulin variable by a defined amplitude *A*_*n*_. We calculate the top Lyapunov exponent Λ_max_ for each inter-kick time as the average of values of 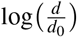from the 1000 relaxation periods. We investigate two such insulin kicking regimes. In the first (Figure 3), we apply kicks of fixed amplitude across all intervals for each inter-kick time value and test two different fixed amplitudes: with A = 5 mU and A = 15 mU. In the second regime (Figure 4), we vary the kick amplitude after every inter-kick interval, pre-drawing 1000 amplitudes from the uniform random distribution *Uniform*(5, 15). To examine how interval length affects the system’s tendency toward chaos, we plot the top Lyapunov exponent Λ_max_ as a function of inter-kick time for plasma insulin under all of the cases described above. Figures 3a and 3b show this comparison for fixed kicking amplitudes (A = 5 mU and A = 15 mU, respectively) under a strictly periodic kicking regime, in which inter-kick interval lengths are fixed across all time points.

**Figure 3.**
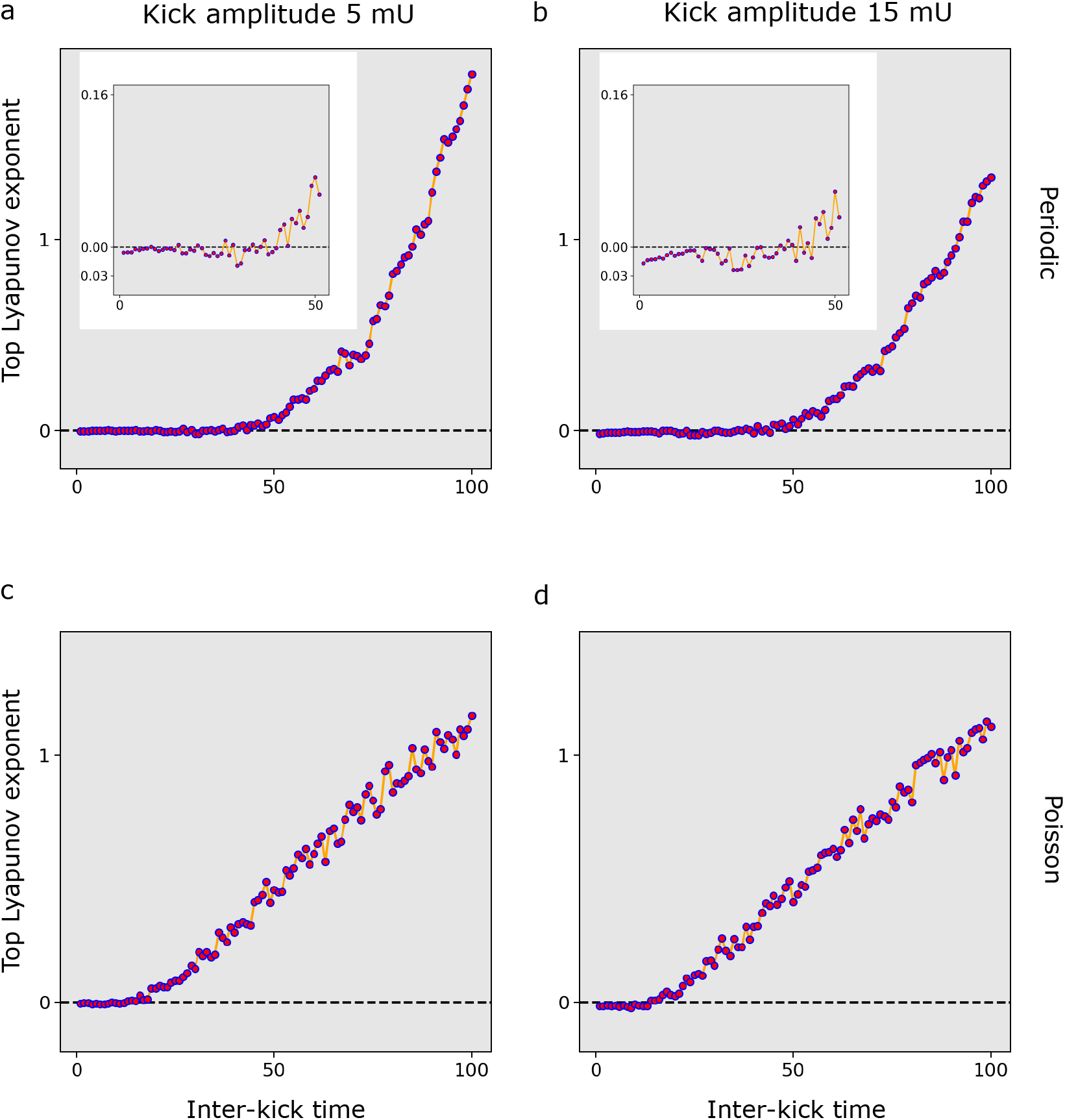
Top Lyapunov exponent Λ_max_ for plasma insulin under constant amplitude pulsatile kicks. The inter-kick time are measured in minutes. The inter-kick times are at periodic intervals in **(a-b)** and drawn independently from an Exponential distribution in **(c-d)**. The first and second column represent kick amplitudes of 5 mU **(a,c)** and 15 mU **(b,d)** respectively.

**Figure 4.**
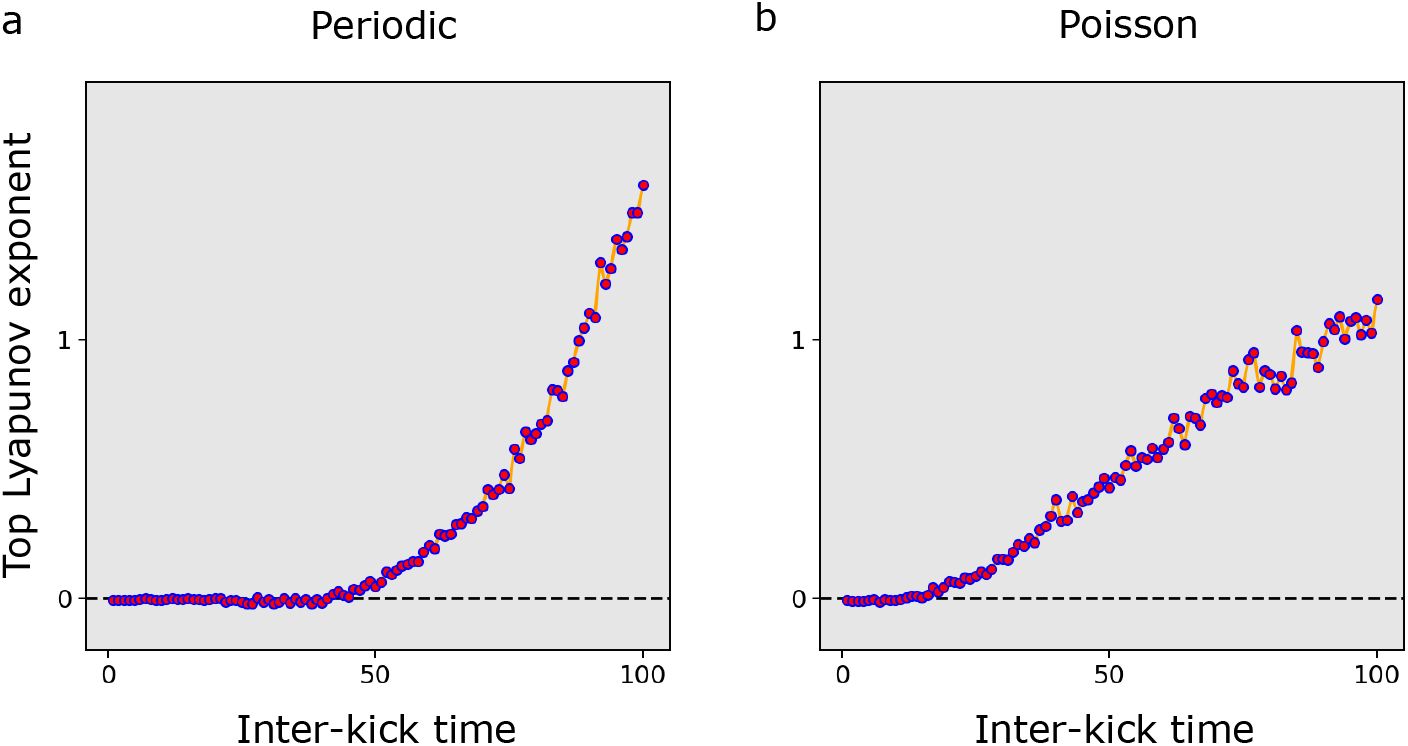
Top Lyapunov exponents for plasma insulin under randomly drawn kick amplitudes with pulsatile kicks. Top Lyapunov exponents Λ_max_ for kick amplitudes drawn from *Uniform*(5, 15) for external pulsatile kicks to plasma insulin. The inter-kick time are measured in minutes. **(a)** Inter-kick times at periodic intervals. **(b)** Inter-kick times drawn independently from an Exponential distribution.

Figures 3c and 3d show the same kick amplitudes applied at varying intervals, with the inter-kick times drawn randomly from an exponential distribution. We then perform a similar comparison under a regime of randomly drawn insulin kick amplitudes, plotting the top Lyapunov exponent against inter-kick time for both periodic (Figure 4a) and randomly drawn (Figure 4b) inter-kick interval lengths.

As inter-kick time increases, we can observe the top Lyapunov exponent becoming positive, signifying the onset of DIU under external pulsatile forcing of plasma insulin. This pattern appears under both constant and randomly varied kicking amplitudes. We also find that longer inter-kick times are required for DIU to emerge when intervals between kicks are periodic than when they are variable. This suggests that the system becomes more prone to chaos with a more realistic pattern of insulin administration.

### 5.2 Kick amplitudes drawn from a Markov process

In Karamched (2021)^2^, we see how DIU can emerge under idealized kicking profiles for glucose. However, glucose consumption in everyday life is far from uniform, meaning that the external forcing amplitudes are not necessarily constant, nor do they strictly follow any known random distribution. Therefore, an alternative approach may be needed to realistically model the type of stochastic glucose kicking profile resulting from ordinary meal consumption or intravenous drips in the ICU.

Generally, meal portions and items vary depending on factors such as hunger intensity, particular food craving, diet, etc. In the ICU, meanwhile, intravenous feeding drips are adjusted depending on the blood glucose level of the patient to maintain glycemic homeostasis. Both of these cases may therefore be reasonably approximated by a kicking profile in which the future kicking amplitude is chosen based on the current glucose amplitude. We can incorporate such a stochastic profile by choosing kick amplitudes based on a Markov process as described in detail by Ross^24^. Let {*X*_*n*_, *n* = 0, 1, 2… }. be a stochastic process in which a state can take a finite or countable number of values. By *X*_*n*_ = *i*, we mean the system is in state *i* at time *n. P*_*i j*_ denotes the probability of entering state *j* from state *i* in a single time step. Here *P*_*i j*_ is only dependent on the state at time *n* and not on any past state of the system. Thus:

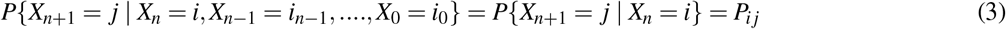

for all states *i*_0_, *i*_1_, ……, *i*_*n−*1_, *i, j* and for all *n ≥* 0. As *P*_*i j*_ denotes the probability of transitioning from *i* to *j* in the next time step, and the system must transition to some state (although it may be identical to the current state) we have *P*_*i j*_ *≥* 0 for all *i, j ≥* 0 and 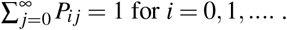The matrix formed with these *P*_*i j*_’s is known as the transition matrix for the Markov process.

Here we approximate the spectrum of possible blood glucose levels with three possible glucose amplitude states: low (*L*), medium (*M*) and high (*H*). The corresponding blood glucose values are 2 mg/dL, 10 mg/dL, and 50 mg/dL, respectively. We choose transition probabilities at random as a simplified proxy for the outcomes of the highly complex real-world processes of meal choice or glucose dosing, resulting in the transition matrix *G*_*l*_:

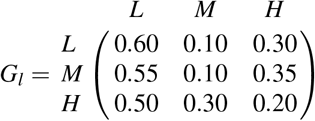

Each element in this matrix shows the probability of transition between two specific glucose states, with rows indicating the present state and columns indicating the subsequent state. For instance, the element in 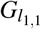 indicates that if the glucose level is in the ‘low’ state at present time step, then it will still be in the ‘low’ state in the next time step with probability 0.60. Similarly, the element in 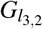 indicates that if the glucose level is ‘high’ at the present time step, then it will transition to the ‘medium’ state at the next time step with probability 0.30. A schematic of this Markov process, including the three plasma glucose states and the associated transition probabilities, is shown in Figure 5.

**Figure 5.**
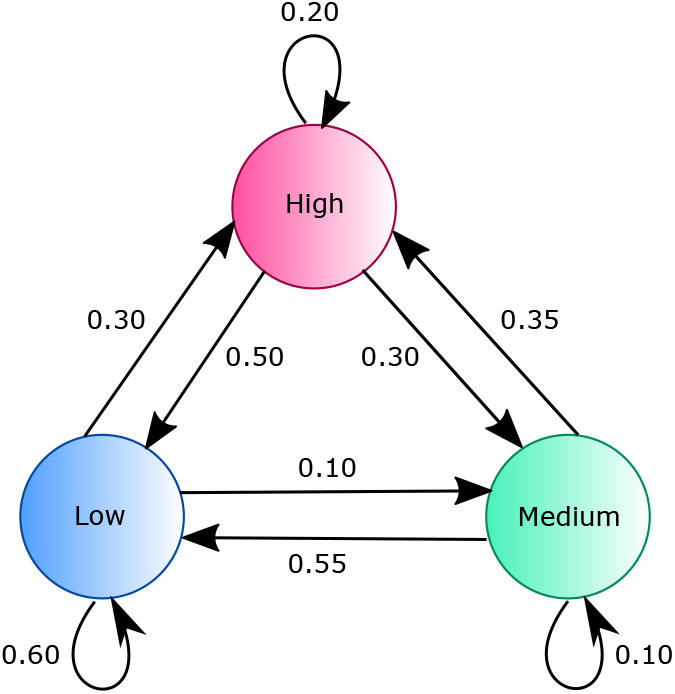
Markov transitions states for glucose. The figure illustrates the Markov states for glucose state transition with respective probabilities.

We draw the external forcing amplitude at each time point in our experiments from the iterated state of this Markov chain, using the numerical values for Low, Medium, and High noted above. To isolate the effect of the time interval between kicks, we vary the inter-kick time but apply the same randomly drawn set of 1000 kick amplitudes to each series.

As seen in Figure 6, under Markov-derived kick amplitudes, the top Lyapunov exponent becomes positive at relatively low inter-kick times (35 minutes or less), marking the onset of DIU in the system. At small values of inter-kick time, the top Lyapunov exponent remains negative, indicating the absence of DIU. We can observe DIU emergence earlier in the Poisson case (roughly 15 minutes) than in the periodic case (roughly 35 minutes), once again suggesting that temporal chaos becomes more likely as the system more closely approximates real-life behavior patterns.

**Figure 6.**
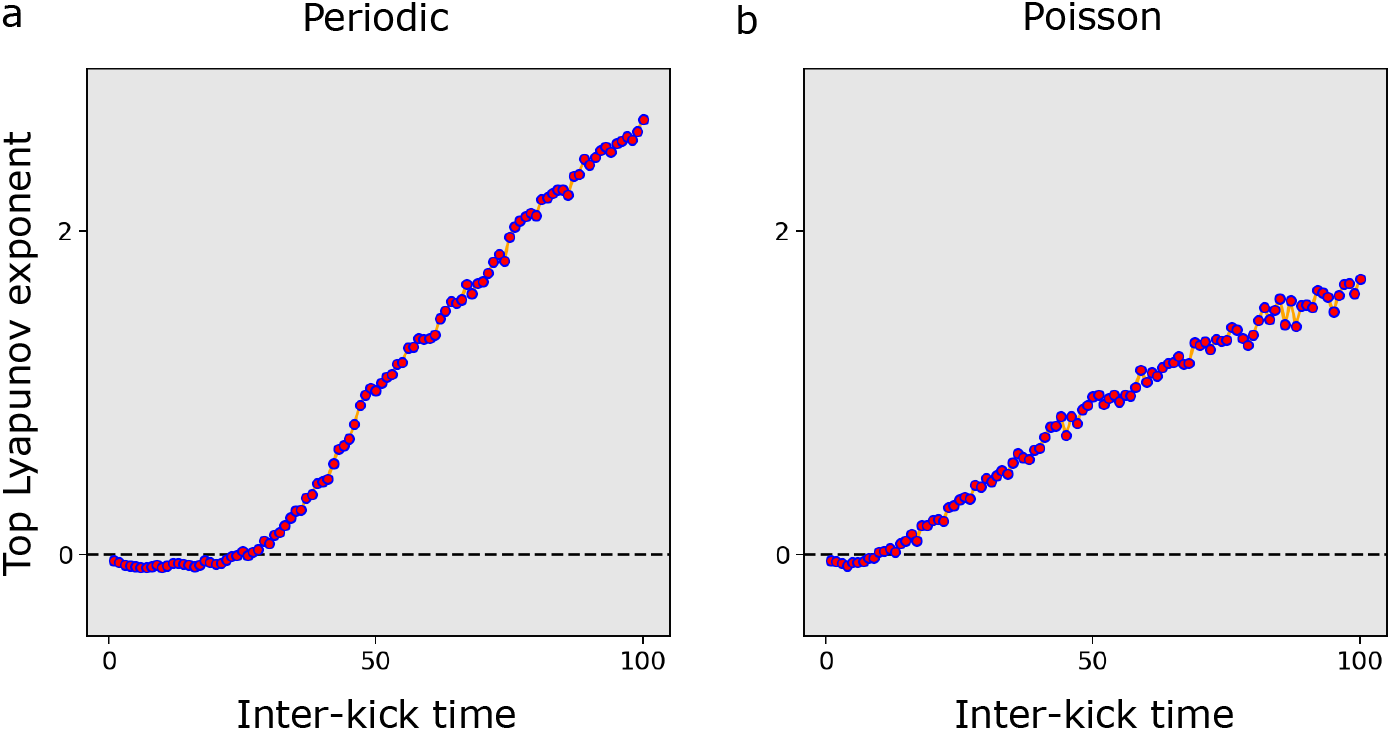
Top Lyapunov exponents for plasma glucose under pulsatile kicks with stochastic amplitudes. Glucose forcing amplitudes are drawn from a Markov process with three states, with magnitudes 2 mg/dL, 10 mg/dL and 50 mg/dL, respectively. Inter-kick time are measured in minutes. **(a)** Top Lyapunov exponents Λ_max_ for periodic inter-kick times. **(b)** Top Lyapunov exponent Λ_max_ for inter-kick times drawn independently from an Exponential distribution.

## 6 Step-wise continuous forcing for glucose

The pulsatile forcing in the previous section is an idealized scenario which assumes that the entire glucose kick enters the patient’s system in a single instantaneous pulse. We aim to make our forcing function much more realistic to what we see in our daily life or the ICU.

So, what changes in the forcing function would help the system more closely mimic realistic scenarios? We consider the case of ICU patients, for whom glucose is administered through intravenous drips over discrete time intervals. We can assume that the glucose forcing amplitude remains constant during these intervals, maintained by the continuous flow of the drip. If needed (i.e., if blood glucose levels become too high or too low), the flow rate will change. We can approximate this scenario using a piece-wise continuous external forcing function.

Let us first carefully note the difference between continuous external forcing functions and the pulsatile kicking described previously in Section 5.2. In contrast to the instantaneous changes in glucose levels modeled in the previous section, here we have continuous glucose forcing. Rather than a ‘kick time’, we now specify a point at which the rate of flow changes, which we call the ‘glucose time change point’. Between any two ‘glucose time change points’, the magnitude of glucose forcing is held constant (Figure 2b). Mathematically, the continuous external forcing is given by:

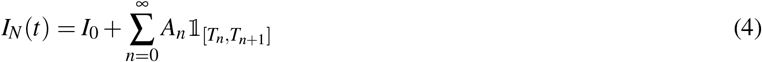

Here *A*_*n*_ denotes the glucose forcing amplitude over [*T*_*n*_, *T*_*n*+1_]. We draw each *A*_*n*_ from a Markov process as in Section 5.2. However, the three glucose states now represent three flow rates for an intravenous drip in the ICU context. ‘Low’ stands for a flow rate of 2 mg dL^*−*1^ min^*−*1^, ‘medium’ stands for a flow rate of 10 mg dL^*−*1^ min^*−*1^ and ‘high’ stands for a flow rate of 50 mg dL^*−*1^ min^*−*1^. We retain the state transition probabilities from the previous section.

First, we simulate the case in which glucose change points occur periodically (*T*_*n*_ = *nT* ), then we repeat the simulation using change points drawn from a Poisson distribution. Here we compute the top Lyapunov exponent as a function of *T* . We use the same numerical scheme described for the case of *δ*-kicks, with one difference: Since *δ*-kicks are no longer present, we do not apply a diffeomorphism at each glucose time change point. We start with the initial Markov glucose state at ‘low’ (2 mg dL^*−*1^ min^*−*1^) and set *I*_0_ = 0 as before.

Once again, we observe the presence of positive top Lyapunov exponents at a fairly small time interval, demonstrating that DIU emerges rapidly in our simulated medical setting.

Similar to our other experiments, this temporal chaos appears both when the glucose time change points are fixed (Figure 7a) and when they are Poisson distributed (Figure 7b). These findings further reinforce our conclusion that the glucose-insulin system is more prone to chaotic behaviour the more closely external forcing conditions approximate real-world glycemic management in the ICU.

**Figure 7.**
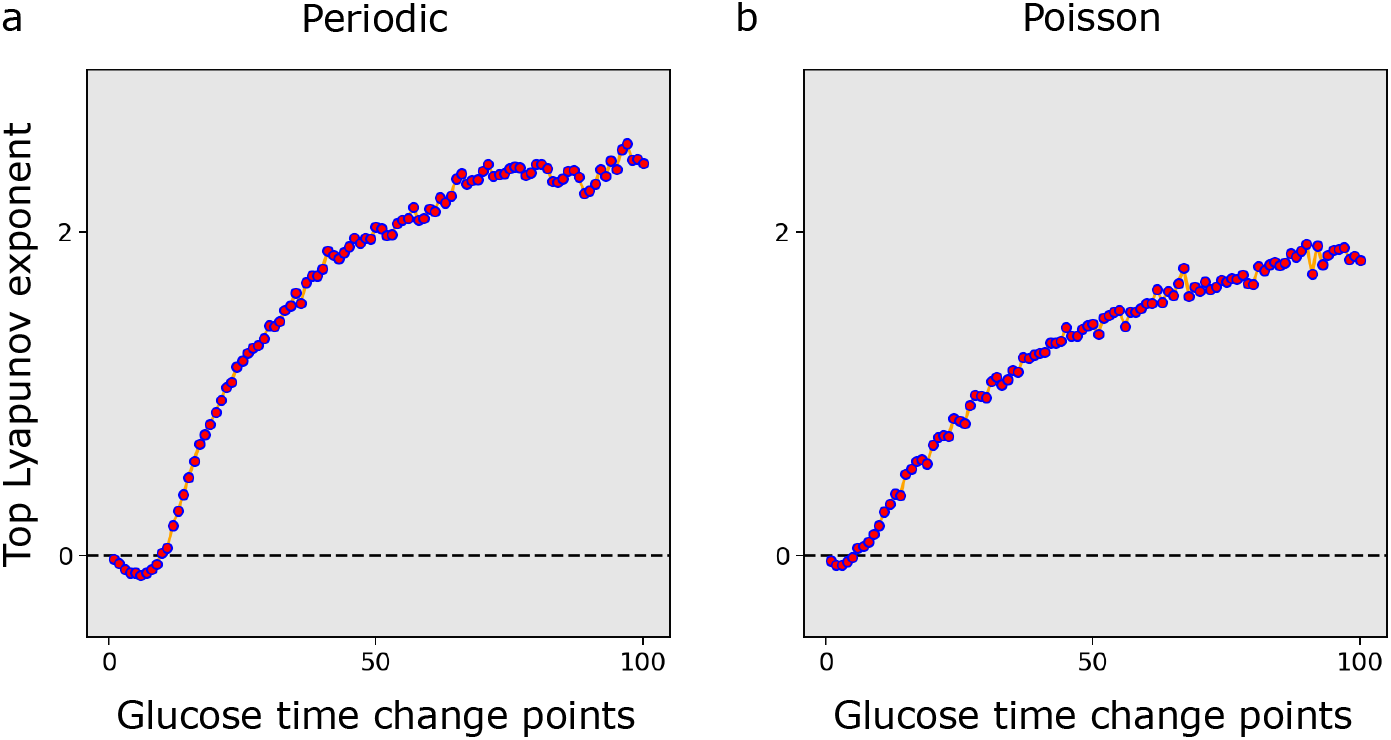
Top Lyapunov exponents for plasma glucose under step-wise continuous forcing. Top Lyapunov exponents Λ_max_ for step-wise external forcing. Amplitudes are drawn from a Markov process with three states, with delivery rate of 2 mg dL^*−*1^ min^*−*1^, 10 mg dL^*−*1^ min^*−*1^ and 50 mg dL^*−*1^ min^*−*1^, respectively. Glucose time change points are measured in minutes. **(a)**Top Lyapunov exponents Λ_max_ for periodic glucose time change points. **(b)** Top Lyapunov exponents Λ_max_ for glucose time change points drawn independently from an Exponential distribution.

## 7 Conclusion and discussions

Through our simulations, we have shown that delay-induced uncertainty is a predictable consequence of the oscillatory dynamics of the glucose-insulin system, and that the patterns of glucose and insulin administration seen in the ICU make the emergence of DIU more likely. First, we demonstrated that pulsatile plasma insulin kicks can drive the emergence of DIU in the Ultradian system. We observed these results under both constant and random kick amplitudes, and for a wide range of time intervals between delta kicks of external insulin forcing. Karamched^2^ reports similar DIU profiles resulting from external glucose forcing under similar kicking profiles. Hence, we have shown the robustness of DIU emergence in the Ultradian model, demonstrating that it is not restricted to any specific state variable.

We further sought to derive glucose kicking profiles that more closely resemble realistic scenarios, drawing kicking amplitudes based on a Markov process to simulate feeding in response to hunger (in everyday meal consumption) or to decreases in measured blood glucose (in the ICU setting). Under these conditions, we show that DIU emerges (as indicated by a positive top Lyapunov exponent) with pulsatile kicks of the glucose state variable. This observation reinforces Karamched’s^2^ prediction that physiological delays should lead to chaotic oscillations in the glucose-insulin system.

Finally, we adjusted our model to account for continuous, dynamically adjusted intravenous administration of glucose as seen in the ICU. We show that this process can be mathematically described as a step-wise function in which the external forcing amplitudes are drawn from the Markov process and remain constant between any two glucose time change points. Our results demonstrate that the appearance of DIU should also be expected in this simulated ICU-like setting. In all of our glucose forcing simulations, we compared kick amplitudes derived from a periodic distribution to those drawn from a Poisson distribution. The former offers a simplified approximation of regular feeding, while the latter more closely approximates the inevitable variation in caloric content between meals or glucose supplements in the real world. We consistently observed DIU at smaller inter-kick intervals in the Poisson case than the periodic case. Natural variation in everyday feeding or medical glucose administration may thus amplify the chaotic tendencies of the glucose-insulin system.

Taken together, our findings suggest that delay-driven chaotic behaviour in mathematical models of glucose-insulin homeostasis is not simply an artifact of simplifying assumptions; in fact, the propensity toward chaos increases as simulations more closely emulate real-world scenarios. We believe that identifying the presence of chaos in the human physiological system and exploring the conditions which lead to such dynamic behaviour will lead to improvements in clinical practice. For example, while artificial intelligence (AI) has advanced medical care and research in numerous ways, our results offer cause for skepticism of purely AI-driven approaches to glycemic management, given the apparent propensity of the underlying system toward chaotic behaviour.

We have designed our experiments to simulate ICU-type conditions and provide numerical evidence that DIU arises in this modeled setting. However, empirical validation will require data from ICU patients, which would allow us to develop external forcing signaling functions that correspond with greater precision to clinical conditions, enhancing our understanding of glycemic management in the ICU setting.

Interestingly, the presence of DIU may not always be a bane to medical management of human physiological systems. Instead, its effects will depend on the clinical setting and the goal we want to achieve. In the ICU, the relevant aim is to maintain glucose homeostasis. Here, the precise temporal evolution of glucose levels may not be important, provided that they remain within the acceptable homeostatic range. Understanding the dynamics of DIU may help clinicians properly tune nutritional input to control the range of glucose-insulin oscillations. On the other hand, DIU may be more problematic when the precise temporal evolution of plasma glucose is clinically relevant (e.g., type-2 diabetes mellitus^9^).

We will close with some questions on which we want to work in the future.

1. How can we demonstrate the presence of DIU in actual ICU patient data to validate our model?
2. Can we translate our findings on DIU in the glucose-insulin system to the many other human physiological systems displaying dynamical features, such as the cardiovascular^25^, pulmonary and respiratory^26,27^, and nervous^28–30^systems? If so, can we apply this understanding to improve clinical practice?
3. How can we account for the fact that the dynamics of glucose-insulin state transitions in living patients can shift over time, rather than remaining fixed as in our simulations?
4. Can our understanding of DIU in the glucose-insulin system help improve type-2 diabetes self-management?
5. Can chaotic behaviors in human homeostatic system arise from other physiological mechanisms besides delay? If so, how can we identify them?

## Code availability

The codes can be found here - https://github.com/Deepgh/Glucose_Insulin_System

## Acknowledgment

We would like to thank William Ott for continuous helpful research discussion and Frederick Naumann for reading and writing help with this paper.

---

**Table 1.** Full list of parameters for the Ultradian glucose-insulin model^22^. Note: IIGU - Insulin-Independent Glucose Utilization and IDGU - Insulin-Dependent Glucose Utilization.

| Ultradian model parameters |  |  |
| --- | --- | --- |
| Name | Nominal Value | Meaning |
| $V_p$ | 3 L | plasma volume |
| $V_i$ | 11 L | interstitial volume |
| $V_g$ | 10 L | glucose space |
| $E$ | $0.2 \text{ L min}^{-1}$ | exchange rate for insulin between remote and plasma compartments |
| $t_p$ | 6 min | time constant for plasma insulin degradation (via kidney and liver filtering) |
| $t_i$ | 100 min | time constant for remote insulin degradation (via muscle and adipose tissue) |
| $t_d$ | 13 min | delay between plasma insulin and glucose production |
| $R_m$ | $209 \text{ mU min}^{-1}$ | linear constant affecting insulin secretion |
| $a_1$ | 6.6 | exponential constant affecting insulin secretion |
| $C_1$ | $300 \text{ mg L}^{-1}$ | exponential constant affecting insulin secretion |
| $C_2$ | $144 \text{ mg L}^{-1}$ | exponential constant affecting IIGU |
| $C_3$ | $100 \text{ mg L}^{-1}$ | linear constant affecting IDGU |
| $C_4$ | $80 \text{ mU L}^{-1}$ | factor affecting IDGU |
| $C_5$ | $26 \text{ mU L}^{-1}$ | exponential constant affecting IDGU |
| $U_b$ | $72 \text{ mg min}^{-1}$ | linear constant affecting IIGU |
| $U_0$ | $4 \text{ mg min}^{-1}$ | linear constant affecting IDGU |
| $U_m$ | $94 \text{ mg min}^{-1}$ | linear constant affecting IDGU |
| $R_g$ | $180 \text{ mg min}^{-1}$ | linear constant affecting IDGU |
| $\alpha$ | 7.5 | exponential constant affecting IDGU |
| $\beta$ | 1.772 | exponent affecting IDGU |

## References

1. Glass, L., Beuter, A. & Larocque, D. Time delays, oscillations, and chaos in physiological control systems. Math. Biosci. 90, 111–125, 10.1016/0025-5564(88)90060-0 (1988).

2. Karamched, B., Hripcsak, G., Albers, D. & Ott, W. Delay-induced uncertainty for a paradigmatic glucose–insulin model. Chaos: An Interdiscip. J. Nonlinear Sci. 31 (2021).

3. Taylor, B. et al. Efficacy and safety of an insulin infusion protocol in a surgical icu. J. Am. Coll. Surg. 202, 1–9, 10.1016/j.jamcollsurg.2005.09.015 (2006).

4. Van den Berghe, G. Beyond diabetes: Saving lives with insulin in the icu. Int. J. Obes. 26, S3–S8, 10.1038/sj.ijo.0802171 (2002).

5. Keener, J. & Sneyd, J. Mathematical physiology, vol. 8 of Interdisciplinary Applied Mathematics (Springer-Verlag, New York, 1998).

6. Sturis, J., Polonsky, K., Mosekilde, E. & Van Cauter, E. Computer model for mechanisms underlying ultradian oscillations of insulin and glucose. Am. J. Physiol. - Endocrinol. Metab. 260, E801–E809, 10.1152/ajpendo.1991.260.5.E801 (1991).

7. Drozdov, A. & Khanina, H. A model for ultradian oscillations of insulin and glucose. Math. Comput. Model. 22, 23–38, 10.1016/0895-7177(95)00108-E (1995).

8. Tolić, I., Mosekilde, E. & Sturis, J. Modeling the insulin-glucose feedback system: The significance of pulsatile insulin secretion. J. Theor. Biol. 207, 361–375, 10.1006/jtbi.2000.2180 (2000).

9. Karamched, B. R., Hripcsak, G., Leibel, R. L., Albers, D. & Ott, W. Delay-induced uncertainty in the glucose-insulin system: Pathogenicity for obesity and type-2 diabetes mellitus. Front. Physiol. 13, 10.3389/fphys.2022.936101 (2022).

10. Zaslavsky, G. The simplest case of a strange attractor. Phys. Lett. A 69, 145–147, 10.1016/0375-9601(78)90195-0 (1978/79).

11. Lin, K. & Young, L.-S. Shear-induced chaos. Nonlinearity 21, 899–922, 10.1088/0951-7715/21/5/002 (2008).

12. Wang, Q. & Young, L.-S. Strange attractors with one direction of instability. Comm. Math. Phys. 218, 1–97, 10.1007/s002200100379 (2001).

13. Wang, Q. & Young, L.-S. Toward a theory of rank one attractors. Ann. Math. (2) 167, 349–480, 10.4007/annals.2008.167.349 (2008).

14. Wang, Q. & Young, L.-S. Dynamical profile of a class of rank-one attractors. Ergod. Theory Dynam. Syst. 33, 1221–1264, 10.1017/S014338571200020X (2013).

15. Ott, W. & Stenlund, M. From limit cycles to strange attractors. Comm. Math. Phys. 296, 215–249, 10.1007/s00220-010-0994-y (2010).

16. Wang, Q. & Young, L.-S. Strange attractors in periodically-kicked limit cycles and Hopf bifurcations. Comm. Math. Phys.240, 509–529, 10.1007/s00220-003-0902-9 (2003).

17. Bergman, R. Origins and history of the minimal model of glucose regulation. Front. Endocrinol. 11, 10.3389/fendo.2020.583016 (2021).

18. Mari, A., Tura, A., Grespan, E. & Bizzotto, R. Mathematical modeling for the physiological and clinical investigation of glucose homeostasis and diabetes. Front. Physiol. 11, 10.3389/fphys.2020.575789 (2020).

19. Hann, C. et al. Integral-based parameter identification for long-term dynamic verification of a glucose-insulin system model. Comput. Methods Programs Biomed. 77, 259–270, 10.1016/j.cmpb.2004.10.006 (2005).

20. Kraegen, E. & Chisholm, D. Insulin responses to varying profiles of subcutaneous insulin infusion: kinetic modelling studies. Diabetologia 26, 208–213, 10.1007/BF00252409 (1984).

21. Wang, H., Li, J. & kuang, Y. Mathematical modeling and qualitative analysis of insulin therapies. Math. Biosci. 210, 17–33, 10.1016/j.mbs.2007.05.008 (2007).

22. Albers, D. et al. Personalized glucose forecasting for type 2 diabetes using data assimilation. PLoS Comput. Biol. 13, 10.1371/journal.pcbi.1005232 (2017).

23. Wilkinson, A. What are lyapunov exponents, and why are they interesting? Bull. Am. Math. Soc. 54, 79–105, 10.1090/bull/1552 (2017).

24. Ross, S. M. Introduction to probability models (Elsevier Academic Press, Amsterdam, 2014), eleventh edn.

25. Christini, D. & Glass, L. Introduction: Mapping and control of complex cardiac arrhythmias. Chaos: An Interdiscip. J. Nonlinear Sci. 12, 732–739, 10.1063/1.1504061 (2002).

26. Mackey, M. & Glass, L. Oscillation and chaos in physiological control systems. Science 197, 287–289, 10.1126/science.267326 (1977).

27. Sottile, P., Albers, D., Higgins, C., Mckeehan, J. & Moss, M. The association between ventilator dyssynchrony, delivered tidal volume, and sedation using a novel automated ventilator dyssynchrony detection algorithm. Critical care medicine 46, e151, 10.1097/CCM.0000000000002849 (2018).

28. Stroh, J., Bennett, T., Kheyfets, V. & Albers, D. Estimating intracranial pressure via low-dimensional models: toward a practical tool for clinical decision support at multi-hour timescales. bioRxiv (2020).

29. Claassen, J. et al. Nonconvulsive seizures after subarachnoid hemorrhage: multimodal detection and outcomes. Annals Neurol. 74, 53–64, 10.1002/ana.23859 (2013).

30. Hodgkin, A. & Huxley, A. A quantitative description of membrane current and its application to conduction and excitation in nerve. The J. Physiol. 117, 500, 10.1113/jphysiol.1952.sp004764 (1952).

